# How to measure functional connectivity using resting-state fMRI? A comprehensive empirical exploration of different connectivity metrics

**DOI:** 10.1101/2024.03.18.585458

**Authors:** Lukas Roell, Stephan Wunderlich, David Roell, Florian Raabe, Elias Wagner, Zhuanghua Shi, Andrea Schmitt, Peter Falkai, Sophia Stoecklein, Daniel Keeser

## Abstract

**Background:** Functional connectivity in the context of functional magnetic resonance imaging is typically quantified by Pearsońs or partial correlation between regional time series of the blood oxygenation level dependent signal. However, a recent interdisciplinary methodological work proposes more than 230 different metrics to measure similarity between different types of time series.

**Objective:** Hence, we systematically evaluated how the results of typical research approaches in functional neuroimaging vary depending on the functional connectivity metric of choice. We further explored which metrics most accurately detect neural decline induced by age and malignant brain tumors, aiming to initiate a debate on how best assessing brain connectivity in functional neuroimaging research.

**Methods:** We addressed both research questions using four independent neuroimaging datasets, comprising multimodal data from a total of 1187 individuals. We analyzed resting-state functional sequences to calculate functional connectivity using 20 representative metrics from four distinct mathematical domains. We further used T1- and T2-weighted images to compute regional brain volumes, diffusion-weighted imaging data to build structural connectomes, and pseudo-continuous arterial spin labeling to measure regional brain perfusion.

**Results:** First, our results demonstrate that the results of typical functional neuroimaging approaches differ fundamentally depending on the functional connectivity metric of choice. Second, we show that correlational and distance metrics are most appropriate to cover neural decline induced by age. In this context, partial correlation performs worse than other correlational metrics. Third, our findings suggest that the FC metric of choice depends on the utilized scanning parameters, the regions of interest, and the individual investigated. Lastly, beyond the major objective of this study, we provide evidence in favor of brain perfusion measured via pseudo-continuous arterial spin labeling as a robust neural entity mirroring age-related neural and cognitive decline.

**Conclusion:** Our empirical evaluation supports a recent theoretical functional connectivity framework. Future functional imaging studies need to comprehensively define the study-specific theoretical property of interest, the methodological property to assess the theoretical property, and the confounding property that may bias the conclusions.

## Introduction

Over the past twenty years, the use of connectivity-based methods has played a leading role in characterizing of both the normal brain organization and alterations due to various brain disorders^1^. In the context of resting-state functional magnetic resonance imaging (rs-fMRI), functional connectivity (FC) reflects the statistical interdependence of the blood-oxygenation-level-dependent (BOLD) signal of two or more brain regions during rest^2–4^. Given an appropriate denoising strategy, the BOLD signal results from changes in cerebral blood flow, volume and oxygenation and is interpreted as an indirect measure of neural activity^4, 5^. Hence, FC refers to the degree of similarity of neural activation between different brain regions^6, 7^. The higher the statistical similarity between the neural activity of different brain regions is, the stronger both regions are assumed to be functionally connected^2, 3^.

Studies that utilize FC as an outcome of interest often apply one of the following approaches: First, based on typically occurring FC patterns, the human brain is divided in several functional brain networks such as the default-mode network^8, 9^. Second, these brain networks can also be regarded as macroscale gradients that distribute along different spatial axes^10^ and represent developmental markers from adolescence to adulthood^11^. Third, many studies attempt to relate certain FC patterns to behavioral domains such as cognitive functioning, aiming to clarify the contribution of certain brain regions to observable human behavior. For instance, FC between the hippocampus and the middle frontal gyrus has been associated with multiple cognitive processes such as working memory^12^. Fourth, to identify robust biomarkers of pathological conditions, many approaches examine alterations in certain FC patterns by comparing clinical samples to matched healthy controls^13–15^. For example, aberrant FC between certain brain networks has been demonstrated in multiple neurological^16^ and psychiatric conditions^13^.

Importantly, most studies that follow one of the four approaches use Pearsońs correlation or partial correlation to quantify FC. Since the correlation coefficient only covers the linear relationship between two BOLD time series, other metrics aiming to assess FC in rs-fMRI research have been proposed in recent years^17–23^. Thereby, exploratory evidence demonstrates that the type of FC metric affects the resulting functional brain configuration regarding the number and size of the extracted brain networks and the regions assigned to each network^17^. A recent interdisciplinary work assigns 237 metrics that quantify similarity between different types of time series to the following six categories: basic measures, distance measures, spectral measures, information-theoretic measures, causal measures, and miscellaneous measures^24^. These similarity metrics are computed based on various distinct approaches and thus differ in their mathematical properties such as dis-/similarity, directionality, directness, domain, or linearity^24^.

With regard to the vast amount of applicable and mathematically different FC metrics, it remains unknown if the abovementioned common approaches in FC research provide stable results independent of the actual FC metric of choice. Therefore, in the first part, we examine the impact of different FC metrics on several common FC-based outcomes in rs-fMRI research. In particular, we assess how FC strength within the default-mode network depends on the actual FC metric of choice. We further inspect the composition of macroscale gradients across different FC metrics. Moreover, we investigate if the association between hippocampal-frontal FC and cognitive functioning remains stable across all FC metrics. Finally, we study if FC between certain brain networks in patients with schizophrenia differs from healthy controls independent of the chosen FC metric.

Irrespective of potential variations across FC metrics, the abovementioned and widely used statistical definition of FC fails to differentiate between the theoretical property of interest and the methodological approach to assess this property^25^. In other words: In many cases, it remains unclear what kind of biological interaction between two brain regions (theoretical property) is covered, when computing the purely statistical correlation between of two BOLD time series (methodological approach). Nowadays, FC has become an inherent entity by itself, although its biological meaning is often not clarified sufficiently^25^. This issue is of particular interest in clinically heterogeneous populations without strong and obvious neurological indications such as psychiatric samples^26^. Hence, the multimodal examination of factors that deteriorate FC patterns in a biologically plausible manner is required^26^ in order to investigate if Pearsońs or partial correlation are the most appropriate metrics to measure connectivity in the human brain.

One such factor is aging, as multiple large-scale examinations have demonstrated a substantial decline of grey and white matter tissue and alterations of brain activation throughout life^27–29^. Another promising factor of interest is a severe neurological condition such as a malignant tumor. Malignant high-grade brain tumors are more likely to invade the surrounding healthy brain tissue, leading to disruptions in white matter integrity and brain function^30, 31^.

Hence, in the second part, we aim to determine which FC metric most accurately captures neural deteriorations and associated cognitive impairments induced by strong biological factors. Accordingly, based on two independent and multimodal large-scale MRI datasets, we evaluate which FC metric is most suitable to detect age-related neural and cognitive decline in the brain. We additionally explore the sensitivity of different FC metrics with regard to local impairments in white matter integrity induced by malignant tumors on the single-patient level. Through our in-depth multimodal and transdiagnostic investigations, we contribute to a broader field of neuroimaging research to better understand if brain connectivity can be quantified in empirical data using metrics other than Pearsońs or partial correlation.

## Methods

### Study Samples

The current study was based on four independent datasets acquired at different sites. We analyzed behavioral and neuroimaging data from the Mind-Brain-Body dataset from the Max-Planck-Institute Leipzig in Germany (MBB, DOI:10.18112/openneuro.ds000221.v1.0.0)^32^, the Clinical Deep Phenotyping cohort from the Department of Psychiatry and Psychotherapy of the Ludwig-Maximilians-University Hospital Munich in Germany (CDP)^33^, the aging cohort of the Human Connectome Project (HCP-Aging)^34^, and from four patients with malignant tumors from the Brain Tumor Connectomics data from the Ghent University Hospital in Belgium (BTC, DOI:10.18112/openneuro.ds001226.v4.0.0)^35, 36^. The CDP and MBB datasets were used to address the first aim of this study, namely to examine the impact of different FC metrics on several common FC-based outcomes in rs-fMRI research. The HCP-Aging cohort, the MBB dataset, and the BTC data were utilized to achieve the studýs second objective, targeting the sensitivity of different FC metrics regarding biological plausible disruptions of neural connections. Table S1 provides an overview of the sample characteristics. Ages and sexes of the four subjects from the BTC dataset are shown in Figure 5.

### MRI Data Acquisition

The MBB dataset was measured at a Siemens 3T Magnetom Verio scanner with a 32-channel head coil, applying a magnetization-prepared two rapid acquisition gradient echoes (MP2-RAGE) sequence, a T2-weighted sequence, a resting-state functional echo-planar-imaging (EPI) sequence, and a diffusion-weighted imaging (DWI) sequence. A detailed description of the scanning protocols is provided elsewhere^32^. The imaging protocol of the CDP study was based on the HCP protoco^34^. It contained a T1-weighted magnetization-prepared rapid acquisition gradient echo (MP-RAGE) sequence, a T2-weighted sampling perfection with application-optimized contrasts using different flip angle evolution (T2-SPACE), a resting-state EPI sequence and a DWI sequence conducted at a Siemens 3T Magnetom Prisma scanner with a 32-channel head coil (for details see Krčmář et al.^33^). The HCP-Aging protocol comprised a T1-weighted MP-RAGE sequence, T2-weighted SPACE sequence, a resting-state EPI sequence, a DWI sequence, and a pseudo continuous Arterial Spin Labeling (PCASL) sequence acquired at a Siemens 3T Magnetom Prisma scanner with a 32-channel head coil. Details are provided in Harms et al.^34^. For the BTC dataset, a T1-weighted MP-RAGE sequence, a resting-state functional EPI sequence, and a multi-shell High Angular Resolution Diffusion Imaging (HARDI) sequence were acquired in a Siemens 3T Magnetom Trio MRI scanner using a 32-channel head coil. The entire scanning protocol is described elsewhere^35, 36^. Table S2 provides an overview of the scanning parameters.

### Multimodal MRI Data Processing

T1- and T2-weighted images of the MBB and HCP-Aging datasets were processed using FreeSurfer v7.2^37–39^. DWI sequences of the MBB, HCP-Aging, and BTC datasets were processed using functions from MRtrix3 v3.0.3^40^, FSL v6.0^41, 42^, Freesurfer v7.2^37–39^, AFNI v22.1.09^43, 44^ and ANTS v2.3.5^45^. PCASL images of the HCP-Aging dataset were processed using Oxford_ASL from FSL v6.0^46^. fMRIPrep v22.1.1^47^ was utilized to preprocess the resting-state functional MRI images of all datasets. After removal of the first ten volumes and subsequent smoothing (FWHM = 6mm), the global signal, cerebrospinal fluid signal, white matter signal, and the extracted noise components from Automatic Removal Of Motion Artifacts based on independent component analysis (ICA-AROMA) were regressed from BOLD time series using the *clean_img* function from Nilearn. The denoised BOLD time series were extracted using the *maskers* module from Nilearn. Subject-specific FC was computed for 20 different metrics using the python-based *pyspi* module^24^.

Details on the preprocessing steps performed for each MRI modality and dataset and the applied quality control procedures are described in the supplemental information.

### Selection of FC Metrics

A recent interdisciplinary study proposed 237 metrics from distinct categories to quantify dis-/similarity between different types of time series and released the python package *pyspi* to compute these metrics^24^. We selected 20 metrics from four mathematical categories already used to quantify FC in the context of resting-state fMRI research. We used Pearsońs correlation, partial correlation, Spearmańs rank correlation, Kendalĺs tau, and cross-correlation as **correlational metrics**. As **distance metrics**, Euclidean distance, cityblock distance, cosine distance, constrained dynamic time warping using the Itakura parallelogram, and constrained dynamic time warping using the Sakoe-Chiba band were computed. Distance metrics were multiplied by minus one prior to data analysis to convert them to similarity measures. Coherence magnitude, phase coherence, phase-locking value, phase-slope index, and spectral Granger causality were assessed as **metrics in the frequency domain**, while mutual information with either a gaussian, kernel-based, or Kraskov-Stögbauer-Grassberger density estimation and transfer entropy with either a gaussian or a Kraskov-Stögbauer-Grassberger density estimation served as **metrics from information theory**. The supplemental information provides a detailed description of these metrics with the respective formulas.

### Multimodal Neuroimaging Outcomes and Cognitive Assessments

As outlined previously, the first objective of this study was to investigate the impact of the selected FC metric on several common FC-based approaches in rs-fMRI research. We considered the following four commonly applied approaches: Investigation of default-mode network connectivity, assessment of macroscale gradients, examination of the association between hippocampal-frontal FC and cognition, and identification of FC-based biomarkers for schizophrenia based on a case-control comparison.

FC within the default-mode network was computed for every subject of the MBB dataset using the aforementioned 20 FC metrics. We individually extracted the BOLD time series for each subject of the MBB dataset, focusing on brain regions within the DMN as defined by the Yeo atlas^9^. Subsequently, we computed FC between these DMN regions for all subjects. The mean FC was then calculated for the 20 metrics, z-standardized, and ranked based on their absolute values, providing insights into the strength of DMN connectivity for the different FC metrics. Additionally, macroscale gradients for each FC metric were evaluated through the BrainSpace toolbox^48^ using the MBB dataset. Recently, the computation of macroscale cortical gradients^10, 48^ has been introduced as a novel approach analyzing BOLD time series in rs-fMRI. Gradient analysis redefines brain segmentation by emphasizing smooth transitions between functional regions rather than rigid boundaries. These gradients reflect transitions both between well-defined networks as per Yeo et al.^9^ and within these networks. For the 20 FC metrics, BOLD time series for each brain region defined by the Schaefer Atlas^49^ of each subject were extracted, resulting in 211 x 20 FC matrices. These were averaged across subjects separately for each metric. Subsequently, gradients were computed for these matrices and compared with gradients initially reported by Margulies et al.^10^. Furthermore, average connectivity matrices were generated for the 20 FC metrics proposed by Cliff et al.^24^ to assess the impact of using metrics with distinct mathematical properties on cortical gradients. Moreover, subjects were stratified by age (in different age bins) and gender patterns to explore additional variations in cortical macroscale gradients.

To cover hippocampal-frontal FC in the healthy subjects of the CDP cohort, several seeds from the hippocampus and medial frontal gyrus were defined based on respective subcortical parcellations from FreeSurfer and the cortical Desikan-Killiany-Tourville (DKT) atlas^50^ (supplemental information). We extracted the respective BOLD time series for each subject and computed FC between hippocampal and frontal seeds for every FC metric. The Brief Cognitive Assessment in Schizophrenia (BACS)^51^ was administered to assess global cognition by taking the average of all z-standardized BACS subdomains as a single composite score.

For the case-control comparison between patients with schizophrenia and healthy subjects of the CDP cohort, we extracted BOLD time series from the seven brain networks defined by Yeo et al.^9^ and calculated subject-specific FC between these networks for each FC metric.

Concerning the second aim of this study, addressing the sensitivity of different FC metrics regarding biologically plausible impairments of neural connections, we used the HCP-Aging cohort and the MBB dataset for age-induced effects, and the BTC dataset to study the impact of malignant tumors. Prior to comparing the FC metrics in terms of their ability to capture age- and tumor-related neural decline, we first intended to demonstrate that neural decline was observable in our particular cohorts using other MRI modalities.

With regard to aging effects on neural and cognitive outcomes, we considered T1- and T2-weighted structural MRI data and DWI data, and PCASL data. Structural MRI and DWI were available for the HCP-Aging and MBB datasets, whereas PCASL data was only provided by the HCP-Aging project. Global cognitive functioning in the HCP-Aging cohort was assessed by the NIH toolbox^52^.

We used the outputs from FreeSurfer v7.2 gained from the structural MRI data to summarize cortical and subcortical brain volumes based on subcortical parcellations from FreeSurfer and the DKT atlas. We first extracted the regional volumes of several previously defined hub regions in the brain (Figure 1, precuneus, posterior cingulate gyrus, anterior cingulate gyrus, insula, superior frontal gyrus, the pallidum, putamen, thalamus, hippocampus)^53, 54^. Volumes were z-standardized and averaged to a score indicating the mean volume in these hub regions. We focused on these hub regions because they are characterized by their particularly pronounced connectivity with all other brain regions^53, 54^.

**Figure 1.**
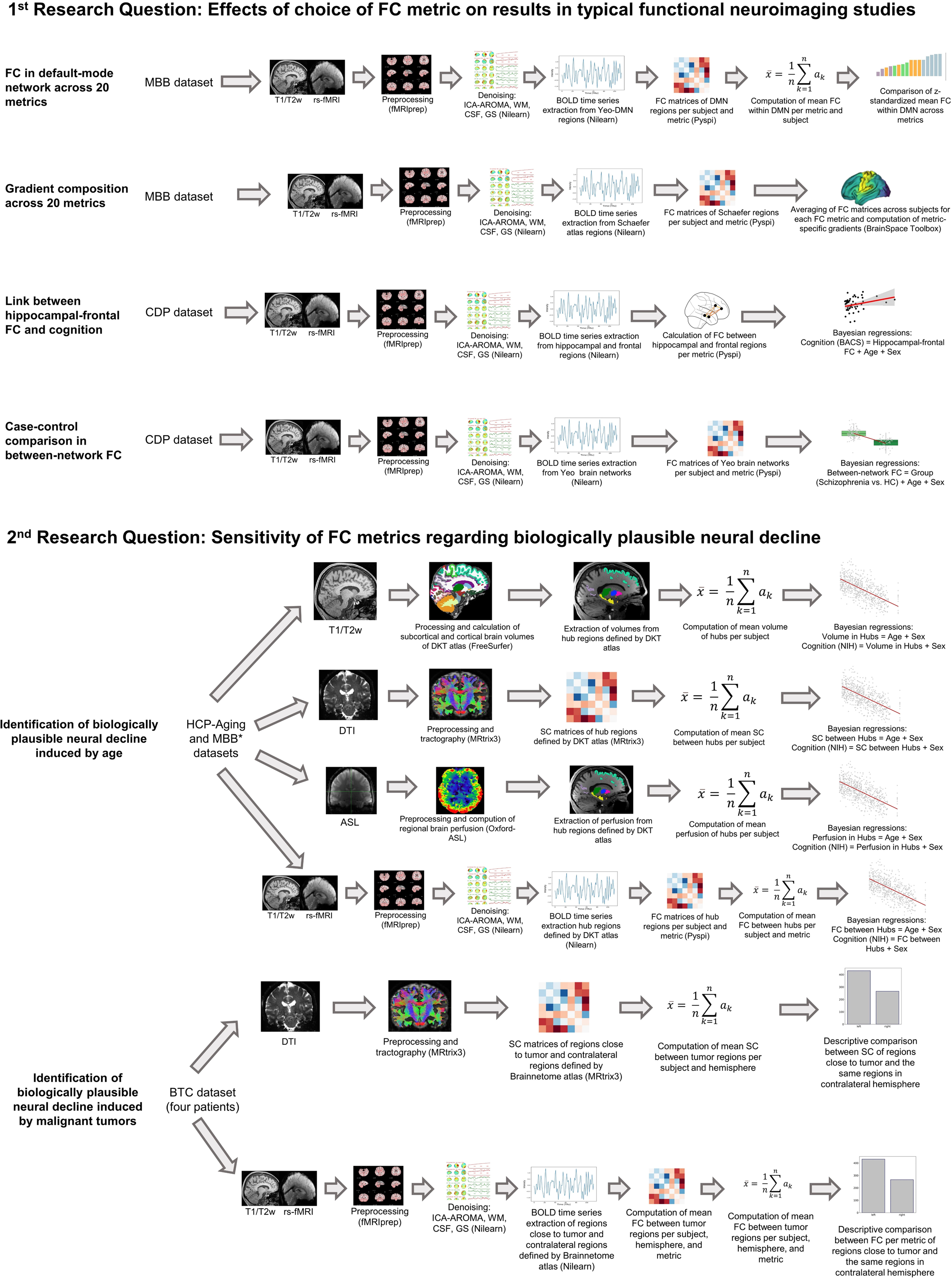
Multimodal processing strategy depending on research question This figure illustrates the processing strategies applied in this study. *, in case of the MBB dataset no PCASL data were acquired and neural outcomes were not linked to cognitive performance. SC, structural connectivity, BACS, Brief Assessment of Cognition in Schizophrenia.

Considering the DWI data, structural connectivity between subcortical and cortical regions defined by subcortical parcellations from FreeSurfer and the DKT atlas was estimated as the number of white matter tracts standardized by the size of the respective regions using the tck2connectome function from MRtrix3^40^. We extracted the structural connectivity values between the hub regions, z-standardized the values, and averaged them to a global structural connectivity score between all hubs.

In case of the PCASL data, Oxford_ASL provided partial volume-corrected measures of the cerebral blood flow for all subcortical and cortical regions specified in the subcortical parcellations from FreeSurfer and the DKT atlas. We extracted the measures of the cerebral blood flow for each hub region, performed a z-standardization, and averaged the values to calculate the mean perfusion across all hubs.

Finally, based on rs-fMRI data, FC between all regions from the subcortical parcellations from FreeSurfer and the DKT atlas was calculated for each FC metric. FC values between all hub regions were extracted, z-standardized, and averaged separately for each FC metric, resulting in one subject-specific value of FC between all hub regions per metric.

We considered DWI and the rs-fMRI data from the BTC dataset to identify biologically plausible neural deteriorations induced by malignant tumors.

Based on the processed DWI data, we computed structural connectivity between all regions from the multimodal Brainnetome atlas^55^ using the tck2connectome function from MRtrix3^40^. For each patient with a malignant tumor, we extracted structural connectivity values between regions close to the specific tumor location and between the same regions from the contralateral hemisphere. Structural connectivity values were z-standardized and averaged to a subject-specific structural connectivity score between regions that were located close to the individual malignant tumor versus a structural connectivity score between the same regions from the contralateral hemisphere. The same strategy was applied for all FC metrics computed from rs-fMRI data.

Figure 1 illustrates the processing strategy and Table S5 provides an overview of all neuroimaging outcomes of interest.

### Statistical Data Analysis

R v4.2.2 was used for statistical data analysis. With regard to the first aim of this study, we evaluated both FC strength within the default-mode network and the composition of macroscale gradients in the MBB dataset descriptively. To investigate if the association between hippocampal-frontal FC and cognitive functioning in the healthy subjects of the CDP cohort remains stable independent of the chosen FC metric, we computed one Bayesian multiple linear regression for each hippocampal-frontal connection and each FC metric with the BACS composite score as dependent variable and hippocampal-frontal FC, age, and sex as predictors. Aiming to evaluate if FC between common brain networks differs between patients with schizophrenia and healthy controls in the CDP cohort independent of the utilized FC metric, we calculated one Bayesian multiple linear regression for each inter-network connection and each FC metric with inter-network FC as dependent variable and group (schizophrenia, healthy controls), age, and sex as predictors.

Regarding the second objective of this study, we explored the association between aging and multimodal neural and cognitive decline in the HCP-Aging and MBB datasets. We used Bayesian multiple linear regressions with either mean volume, mean perfusion, mean structural connectivity, and mean functional connectivity (for all 20 metrics) of the hub regions as dependent variables and age and sex as predictors. Note that we used age as a binary predictor in the case of the MBB dataset (younger adults between 20 and 35 years versus older adults between 55 and 80 years), because only age categories instead of actual ages of participants were published for this dataset. Moreover, we calculated Bayesian multiple linear regressions with the cognitive composite score from the NIH toolbox as dependent variable and either mean volume, mean perfusion, mean structural connectivity, and mean functional connectivity (for all 20 metrics) of the hub regions and sex as predictors. Structural connectivity and metric-specific FC between regions close to the malignant tumor and the same regions from the contralateral hemisphere were compared descriptively.

The main test statistics of interest from the Bayesian multiple linear regression computed with the *brms* package^56^ was Jeffreýs default Bayes Factor (BF_10_), representing a continuous, relative measure of evidence the data is providing for the alternative hypothesis (H_1_: β_z_ ≠ 0) compared to the null hypothesis (H_0_: β_z_ = 0)^57^. A BF_10_ between one and three reflects anecdotal evidence for the alternative hypothesis, between three and ten is considered to be moderate evidence, between ten and 30 is labeled as strong evidence, between 30 and 100 is seen as very strong evidence, and a BF_10_ above 100 is decisive evidence for the alternative hypothesis^58^.

## Results

### Default-mode network connectivity across FC metrics

We compared the absolute connectivity strength for all 20 FC metrics across both hemispheres. Based on the distance metrics, we obtained the highest absolute connectivity strength within the DMN followed by the correlational metrics, whereas the information-theory and frequency metrics showed the lowest. Among the correlational metrics, DMN connectivity was the lowest for partial correlation (Figure S1).

### Composition of macroscale functional gradients across FC metrics

Depending on the FC metric used, gradient patterns vary in their graphical representation, as indicated in Figure S2. Especially, phase coherence, dynamic time warping with an Itakura constraint, spectral Granger causality, and transfer entropy with a Kraskov-Stögbauer-Grassberger density estimation do not show patterns related to the DMN as proposed in Margulies et al.^10^. In contrast, the remaining FC metrics show similar patterns, most dominantly reflected by the correlational metrics. Importantly, the first and second gradients were inverted for some metrics compared to Margulies et al.^10^.

### Association between hippocampal-frontal FC and cognition across FC metrics

Figure 2A illustrates the β-coefficients and BF_10_ of the respective hippocampal-frontal connection extracted from the Bayesian multiple linear regression that assessed the association between hippocampal-frontal FC and cognition using the healthy controls from the CDP cohort. Results indicate that depending on the FC metric of choice a different number and different types of hippocampal-frontal connections show associations with cognitive functioning. For instance, when using Pearsońs correlation, Spearmańs correlation, Kendalĺs tau, Euclidean distance, Cityblock distance, Cosine distance or dynamic time warping with a Sakoe-Chiba band, higher FC between the left parahippocampal gyrus and the left caudal anterior cingulate gyrus is related to better cognitive performance. This finding is not reproduced in the case of partial correlation or any other FC metric, demonstrating that the choice of the FC metric affects the results and interpretations of brain-behavior associations.

**Figure 2.**
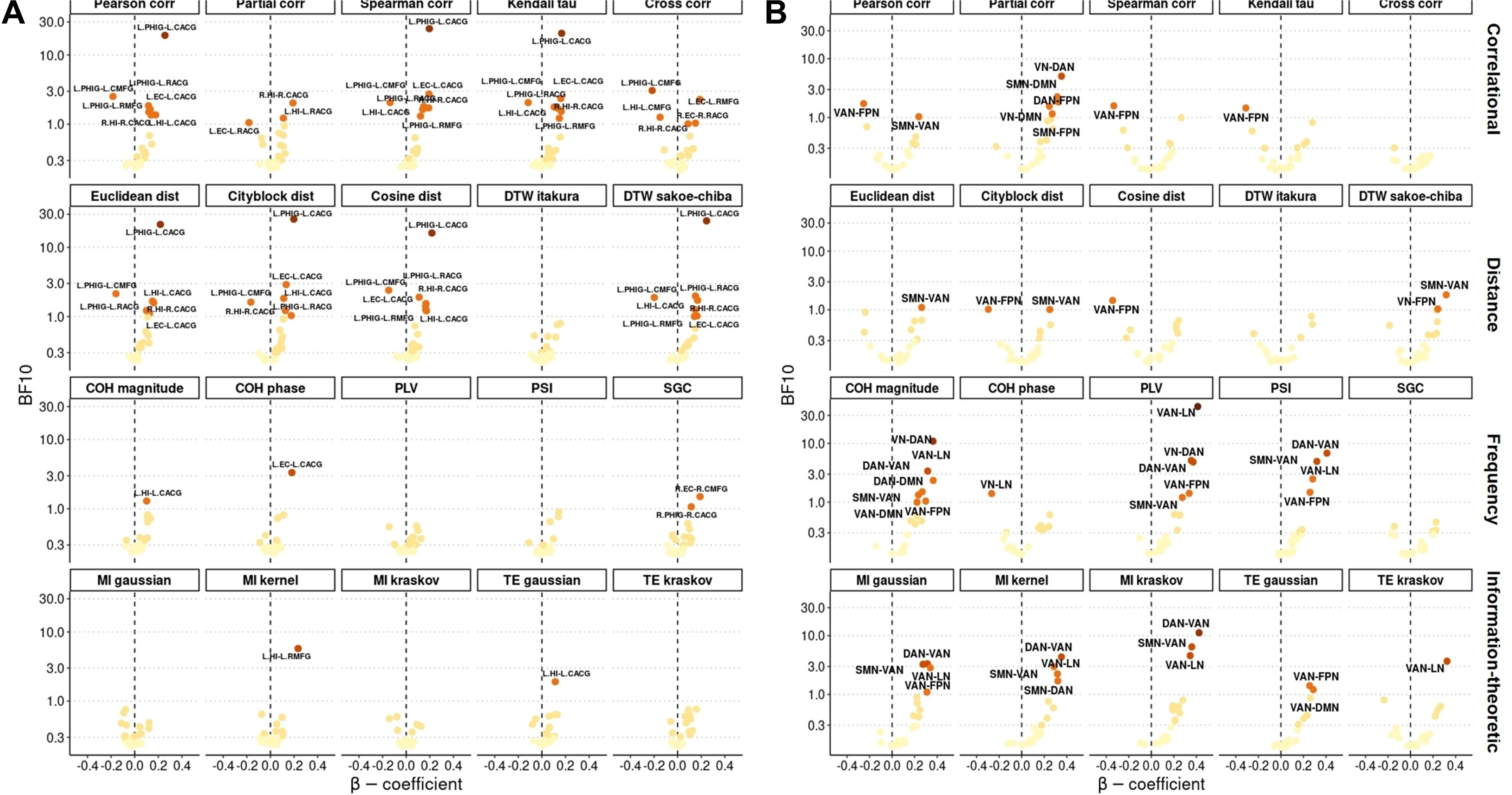
Brain-behavior associations and dysconnectivity patterns between cases and controls for 20 metrics This figure displays the results from Bayesian multiple linear regression analyses, addressing the associations between hippocampal-frontal FC and cognitive functioning **(A)** and the FC differences between patients with schizophrenia and healthy controls **(B)** across 20 FC metrics. Each dot reflects one Bayesian multiple linear regression. **(A)** The β-coefficient of the respective hippocampal-frontal connection is shown on the x-axis and the BF10 of this predictor is illustrated on the y-axis. A positive β indicates a positive association between the particular functional connection and cognition, and a negative β reveals a negative association. **(B)** The β-coefficient of the predictor group (schizophrenia vs. controls) is shown on the x-axis and the BF10 of this predictor is illustrated on the y-axis. A positive β indicates a higher FC between the respective networks in patients with schizophrenia and a negative β reveals lower FC. The higher the BF10, the darker the color of the dots. Functional connections are labelled if the BF10 is higher than one. Pearson corr, Pearsońs correlation; Partial corr, partial correlation; Spearman corr, Spearmańs correlation; Kendall tau, Kendalĺs tau; Cross corr, cross correlation; Euclidean dist, Euclidean distance; Cityblock dist, Cityblock distance; Cosine dist, Cosine distance; DTW itakura, dynamic time warping constrained with Itakura parallelogram; DTW sakoe-chiba; dynamic time warping constrained with Sakoe-Chiba band; COH magnitude, coherence magnitude; COH phase, phase coherence; PLV, phase-locking value; PSI, phase slope index; SGC, spectral Granger causality; MI gaussian, mutual information with gaussian density estimation; MI kernel, mutual information with kernel-based density estimation; MI kraskov, mutual information with Kraskov-Stögbauer-Grassberger density estimation; TE gaussian, transfer entropy with gaussian density estimation; TE kraskov transfer entropy with Kraskov-Stögbauer-Grassberger density estimation; L, left hemisphere; R, right hemisphere; PHIG, parahippocampal gyrus; CMFG, caudal middle frontal gyrus; CACG, caudal anterior cingulate gyrus; RACG, rostral anterior cingulate gyrus; RMFG, rostral middle frontal gyrus; EC, entorhinal cortex; HI, hippocampus; VN, visual network, SMN, somatormotor network; DAN, dorsal attention network; VAN, ventral attention network; LN, limbic network; FPN, fronto-parietal network; DMN, default-mode network.

### Dysconnectivity patterns in schizophrenia across FC metrics

Figure 2B shows the β-coefficients and BF_10_ of the predictor group extracted from the Bayesian multiple linear regression that covered the case-control differences in FC between brain networks based on patients with schizophrenia and healthy controls from the CDP study. Results reveal that depending on the FC metric of choice different numbers and types of dysconnectivity patterns are found. For example, when using partial correlation, coherence magnitude or phase-locking value, patients with schizophrenia reveal a hyperconnectivity between the visual network and the dorsal attention network. At the same time, this is not the case for any other FC metric. Hence, the choice of the FC metric impacts the results of disorder-related dysconnectivity patterns.

### Sensitivity of FC metrics regarding age-related neural and cognitive decline

Figures 3 and 4 visualize the associations between age and multimodal neural outcomes in the hub regions and the test statistics extracted from the respective Bayesian multiple linear regressions based on the HCP-Aging and the MBB cohort. With regard to the HCP-Aging dataset (Figure 3A-D), we found decisive evidence that the older the participants were, the lower their mean volume (Figure 3A), structural connectivity (Figure 3B), and perfusion (Figure 3C) in the hub regions was, on average. The strongest effects were found for volume, followed by perfusion and structural connectivity. These findings demonstrate that multimodal age-related neural decline in the hub regions was present in the HCP-Aging cohort. Considering the rs-fMRI data (Figure 3D), our findings reveal decisive evidence that FC between the hub regions decreases with age, when using Pearsońs correlation, Spearmańs correlation, Kendalĺs tau, cross correlation, Euclidean distance, or cosine distance as FC metric. Correspondingly, we found very strong evidence for this association with Cityblock distance and strong evidence when utilizing Gaussian mutual information. In the case of partial correlation and dynamic time warping with a Sakoe-Chiba band we only found anecdotal evidence, but strong and anecdotal evidence for phase coherence and gaussian transfer entropy towards a positive relation between age and FC in hub regions, respectively. Effect sizes were overall smaller than in the other MRI modalities. With respect to the MBB dataset (Figure 3E-G), we found decisive evidence that elderly, on average, had lower mean volume (Figure 3E) and structural connectivity (Figure 3F) in the hub regions than younger participants. The effect for volume was stronger than the effect for structural connectivity. These findings demonstrate that multimodal age-related neural decline in the hub regions was also observable in the MBB cohort. Considering the rs-fMRI data (Figure 3G), our findings demonstrate decisive evidence that the elderly show lower FC between the hub regions than younger participants, when using cross-correlation and dynamic time warping constrained by an Itakura parallelogram. We also obtained strong evidence for this group difference in the case of Pearsońs correlation, Spearmańs correlation, Kendalĺs tau, and cosine distance, but moderate and anecdotal evidence by Euclidean distance and partial correlation, respectively. Higher FC between the hub regions in elderly was evident when using dynamic time warping with a Sakoe-Chiba band, coherence magnitude, phase coherence, phase-locking value, spectral Granger causality, and both transfer entropy versions. Apart from dynamic time warping constrained by an Itakura parallelogram, effect sizes were smaller than in the other MRI modalities.

**Figure 3.**
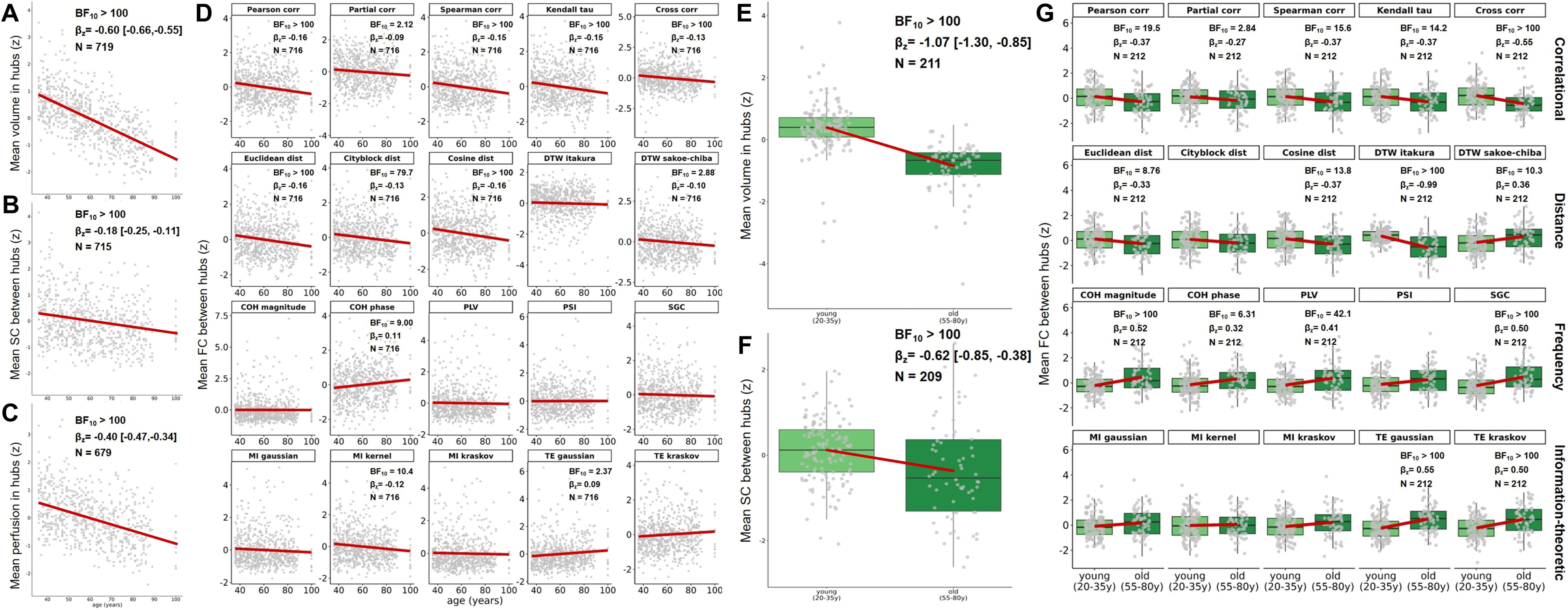
Associations between age and multimodal neural outcomes for the HCP-Aging and MBB cohort This figure depicts the associations between age and mean volume, structural connectivity, perfusion, and functional connectivity in the hub regions, as well as the test statistics of the respective Bayesian multiple linear regressions. Each dot represents a healthy subject from the HCP-Aging (A-D) and the MBB (E-G) cohort. The red regression line reflects the correlation between age and the particular neural outcome. A) Association between age and mean volume in hub regions with age displayed on the x-axis and mean volume on the y-axis. B) Association between age and mean structural connectivity between hub regions with age displayed on the x-axis and mean structural connectivity on the y-axis. C) Association between age and mean perfusion in hub regions with age displayed on the x-axis and mean perfusion on the y-axis. D) Association between age and mean FC between hub regions with age displayed on the x-axis and mean FC on the y-axis for each FC metric. E) Difference in mean volume in hub regions with the age groups displayed on the x-axis and mean volume on the y-axis. F) Difference in mean structural connectivity between hub regions with the age groups displayed on the x-axis and mean structural connectivity on the y-axis. G) Difference in mean FC between hub regions with the age groups displayed on the x-axis and mean FC on the y-axis for each FC metric. Pearson corr, Pearsońs correlation; Partial corr, partial correlation; Spearman corr, Spearmańs correlation; Kendall tau, Kendalĺs tau; Cross corr, cross correlation; Euclidean dist, Euclidean distance; Cityblock dist, Cityblock distance; Cosine dist, Cosine distance; DTW itakura, dynamic time warping constrained with Itakura parallelogram; DTW sakoe-chiba; dynamic time warping constrained with Sakoe-Chiba band; COH magnitude, coherence magnitude; COH phase, phase coherence; PLV, phase-locking value; PSI, phase slope index; SGC, spectral Granger causality; MI gaussian, mutual information with gaussian density estimation; MI kernel, mutual information with kernel-based density estimation; MI kraskov, mutual information with Kraskov-Stögbauer-Grassberger density estimation; TE gaussian, transfer entropy with gaussian density estimation; TE kraskov transfer entropy with Kraskov-Stögbauer-Grassberger density estimation; BF10, Bayes factor of the predictor age; βz, standardized beta coefficient of the predictor age and 95% confidence interval; N, sample size considered in the respective analysis; SC, structural connectivity; FC, functional connectivity.

**Figure 4.**
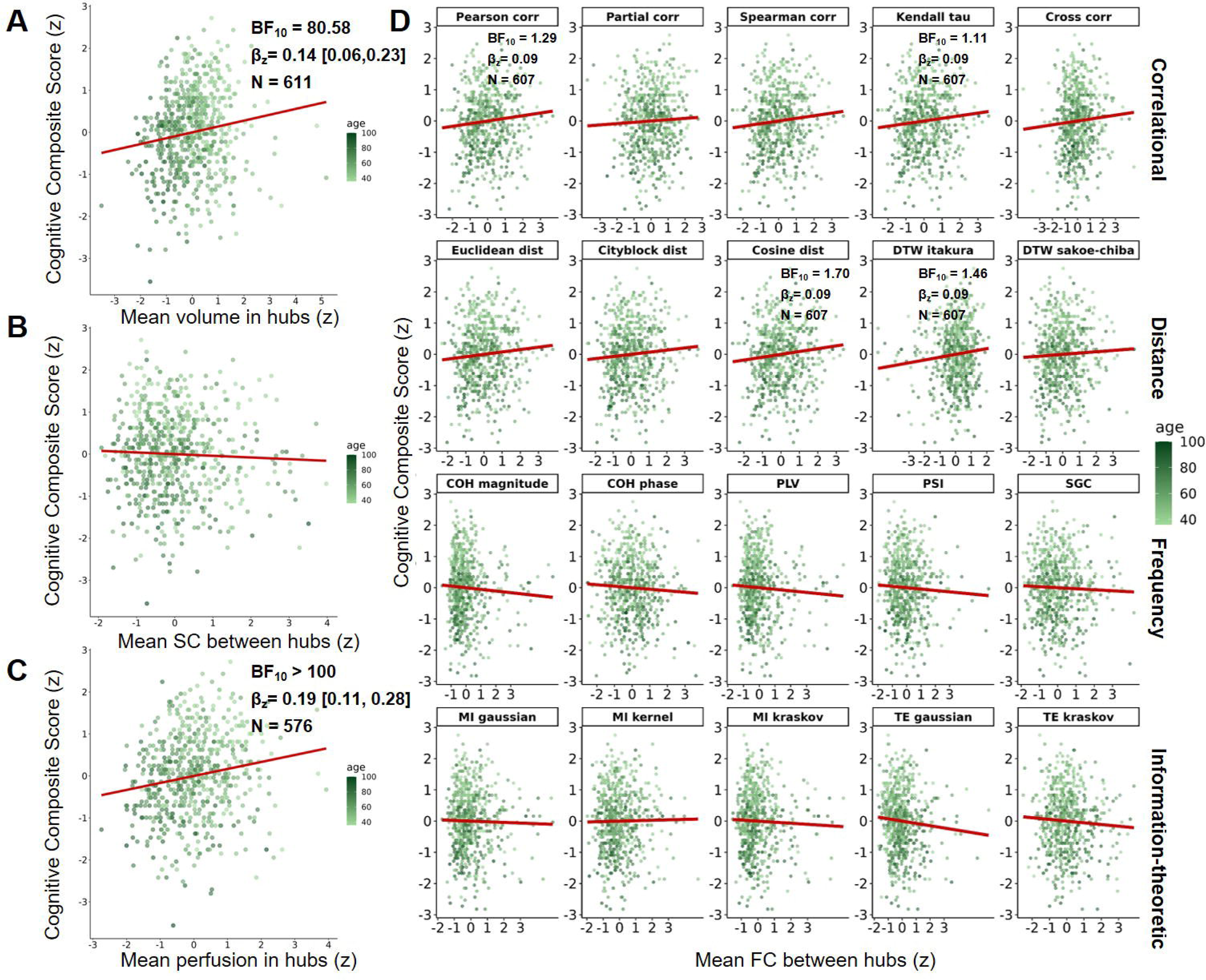
Associations between multimodal neural outcomes and global cognition for the HCP-Aging cohort This figure shows the associations between mean volume, structural connectivity, perfusion, and functional connectivity in the hub regions and the cognitive composite score, as well as the test statistics of the respective Bayesian multiple linear regressions. Each dot represents on healthy subject from the HCP-Aging cohort. The red regression line reflects the correlation between the particular neural outcome and cognitive performance. A) Association between mean volume in hub regions and cognition with mean volume displayed on the x-axis and the cognitive composite score on the y-axis. B) Association between mean structural connectivity between hub regions and cognition with mean structural connectivity displayed on the x-axis and the cognitive composite score on the y-axis. C) Association between mean perfusion in hub regions and cognition with mean perfusion displayed on the x-axis and the cognitive composite score on the y-axis. D) Association between mean FC between hub regions and cognition with mean FC between hub regions displayed on the x-axis and the cognitive composite score on the y-axis. Pearson corr, Pearsońs correlation; Partial corr, partial correlation; Spearman corr, Spearmańs correlation; Kendall tau, Kendalĺs tau; Cross corr, cross correlation; Euclidean dist, Euclidean distance; Cityblock dist, Cityblock distance; Cosine dist, Cosine distance; DTW itakura, dynamic time warping constrained with Itakura parallelogram; DTW sakoe-chiba; dynamic time warping constrained with Sakoe-Chiba band; COH magnitude, coherence magnitude; COH phase, phase coherence; PLV, phase-locking value; PSI, phase slope index; SGC, spectral Granger causality; MI gaussian, mutual information with gaussian density estimation; MI kernel, mutual information with kernel-based density estimation; MI kraskov, mutual information with Kraskov-Stögbauer-Grassberger density estimation; TE gaussian, transfer entropy with gaussian density estimation; TE kraskov transfer entropy with Kraskov-Stögbauer-Grassberger density estimation; BF10, Bayes factor of the predictor age; βz, standardized beta coefficient of the predictor age and 95% confidence interval; N, sample size considered in the respective analysis; SC, structural connectivity; FC, functional connectivity.

**Figure 5.**
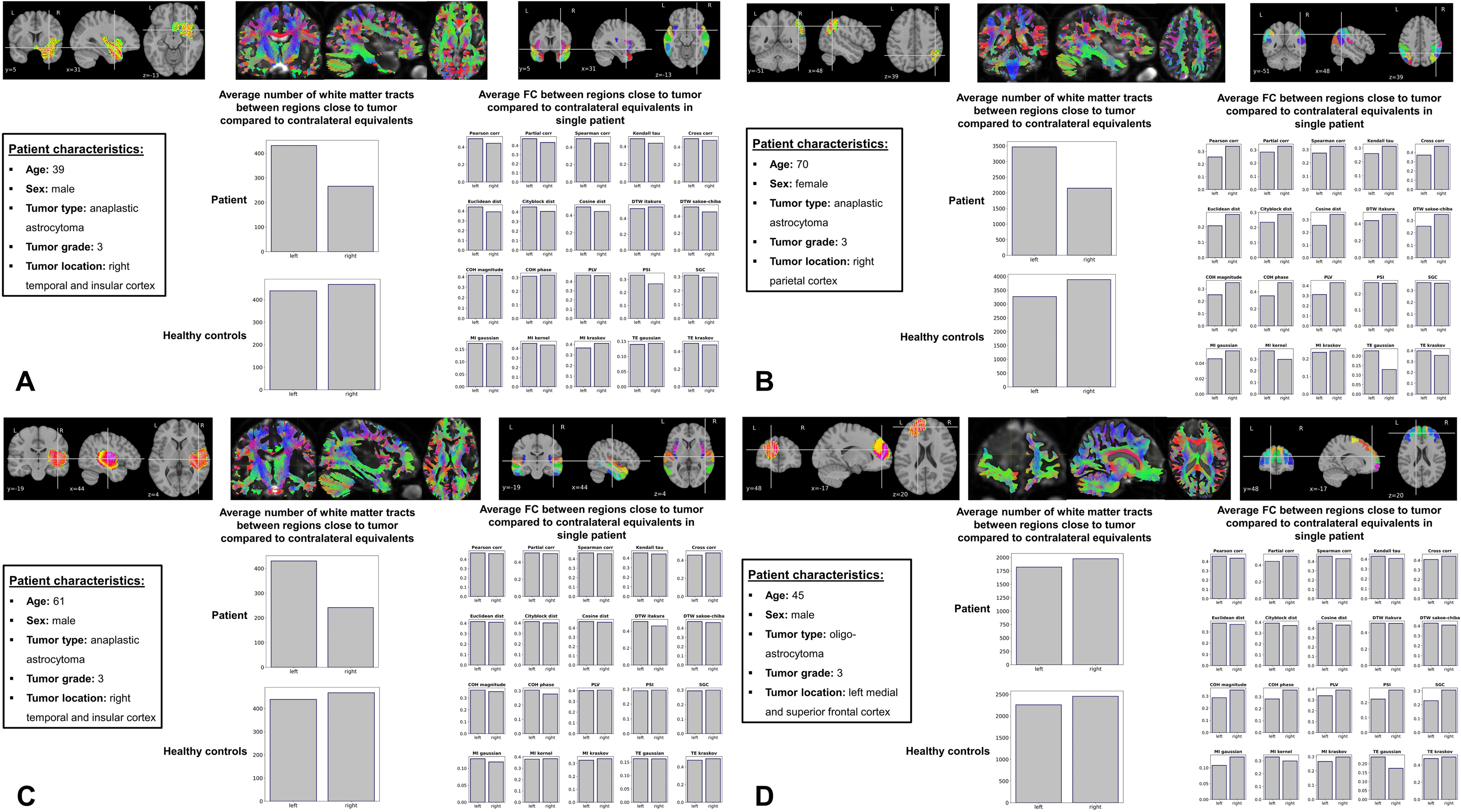
Comparison of white matter tracts and FC between regions close to tumor and contralateral equivalents This figure shows the average number of white matter tracts between regions close to the subject-specific tumor location compared to the same regions in the contralateral hemisphere for four subjects with malignant tumors in the BTC dataset and for respective healthy controls. In each panel (A-D), the tumoŕs location is shown on the top left, the respective tractography on the top middle, and the regions from the Brainnetome atlas used to compute the average number of white matter tracts and FC on the top right. Patient characteristics and the comparison in the average number of white matter tracts and FC between selected regions from both hemispheres are illustrated.

Referring back to the HCP-Aging cohort, we obtained very strong and decisive evidence that larger mean volumes (Figure 4A) and higher mean perfusion (Figure 4C) in the hub regions, respectively, were linked to better cognitive performance in the HCP-Aging cohort. The effect for perfusion was stronger than the effect for volume. We did not encounter such an association for structural connectivity between hub regions (Figure 4B). These findings demonstrate that age-related cognitive decline in the hub regions was present for volumes and perfusion in the HCP-Aging cohort. Regarding the rs-fMRI data (Figure 4D), only anecdotal evidence was observed, suggesting that higher FC between the hub regions was linked to better cognitive performance, when using Pearsońs correlation, Kendalĺs tau, cosine distance, and dynamic time warping constrained with an Itakura parallelogram. Effect sizes were smaller than in structural MRI and PCASL.

### Sensitivity of FC metrics regarding tumor-induced neural decline

Figure 5 displays structural connectivity and FC data from the BTC dataset for four patients with malignant tumors.

Our findings indicate that the average number of white matter tracts between regions close to the subject-specific tumor location was smaller compared to the same regions in the contralateral hemisphere for the first three subjects, whereas this was not the case for the healthy controls. For the first subject (Figure 5A), lower FC between regions close to the subject-specific tumor location compared to the same regions in the contralateral hemisphere was observed, when using Pearsońs correlation, Spearmańs correlation, Kendalĺs tau, Euclidean distance, Cityblock distance, cosine distance, dynamic time warping constrained by a Sakoe-Chiba band, or the phase-slope index as FC metrics. In case of the second subject (Figure 5B), only kernel-based mutual information and both transfer entropy metrics revealed corresponding results. In contrast, correlational and distance metrics, as well as several frequency metrics indicated an effect in the opposite direction. For the third subject (Figure 5C), no FC metric except phase coherence and dynamic time warping constrained by an Itakura parallelogram revealed clear differences in FC between regions close to the subject-specific tumor location, which was smaller compared to the same regions in the contralateral hemisphere. In case of the fourth subject (Figure 5D), the average number of white matter tracts did differ substantially between both hemispheres and showed a similar pattern like the healthy controls. However, based on partial correlation and the frequency measures reduced FC was observed in the ipsilateral hemisphere.

## Discussion

Based on four independent multimodal MRI data sets, our study first explored if FC metrics with distinct mathematical properties lead to different conclusions in the context of typical FC-based research questions. Furthermore, we evaluated whether FC metrics other than Pearsońs or partial correlation are more appropriate for detecting apparent neural deteriorations induced by age or malignant brain tumors.

### Influence of FC Metrics on Typical Research Approaches in Functional Neuroimaging

Our findings suggest that the selected FC metric affects the results and conclusions in several common FC-based research approaches, such as examining FC within the default-mode network, exploring the composition of macroscale gradients, investigating brain-behavior associations, and studying disorder-related dysconnectivity patterns. This lines with previous evidence demonstrating that distinct FC metrics computed within the same dataset lead to substantial differences in the resulting brain network configuration^17^. Hence, the choice of FC metric represents an essential step in studies including FC as a relevant outcome, because the resulting conclusions on the particular research questions will be fundamentally different. With respect to the noticeable mathematical differences of potentially applicable FC metrics, the resulting variation in typical FC-based outcomes is not surprising. In particular, the FC metrics examined in this study differ in terms of their type (similarity or dissimilarity measure or both), directionality (non-directional, unidirectional, or bidirectional), directness (direct or non-direct measure), domain (time or frequency), and linearity assumption (linear or non-linear)^24, 59^. For instance, partial correlation reflects both similarity and dissimilarity between two BOLD time series. It does not assume a direction of the association, considers the influence of other regions, and thus assesses the direct association between the two regions of interest. Moreover, partial correlation captures similarity in the time domain and includes a linearity assumption. The supplemental information provides an overview of the mathematical properties of all applied FC metrics.

### Superiority of Correlational and Distance Metrics in Detecting Age-related Neural Decline

Apart from demonstrating the impact of different FC metrics on the results in common FC-based research approaches, our findings reveal that correlational and distance metrics most accurately capture neural decline. Although generally in line with previous simulation studies that also suggest the superiority of correlational measures in detecting specific alterations in brain networks^60^, we observed that partial correlation only provided anecdotal evidence in favor of age-related neural decline, whereas other correlational and distance metrics indicated more robust associations. In contrast, partial correlation was found to be one of the most accurate FC metrics when tested in simulated BOLD time series^60^ and is generally suggested to be less prone to confounds resulting from other neural entities^25^. Despite these convincing examinations and the underlying reasonable theoretical foundation, our findings suggest that using partial correlation should have some additional consideration. Here, we used a rather global measure of FC as the main outcome by averaging the subject-specific FC values per metric between all hub regions. If a certain brain region is highly connected to many other regions, it is more likely to influence FC between all other regions. When not controlling for its impact on other functional connections, this highly connected region will be weighted stronger than less connected regions in an average FC value used in this study. Given that we demonstrated an age-related decline of white matter tracts between hub regions, a summarizing FC that gives more weight to highly connected regions may be beneficial in covering neural and cognitive deteriorations induced by age. This emphasizes the role of directness as one important property of FC metrics.

However, directness is not the only important property, as FC metrics in the frequency domain and from information theory did not capture age-related neural and cognitive decline properly, despite also covering indirect associations between brain regions like most correlation and distance metrics. In the case of frequency metrics computed from BOLD time series, previous simulation studies raise concerns about their sensitivity regarding alterations in the underlying network architecture^60^, which corresponds to our empirical findings. These concerns also apply to the directional metrics we examined, namely spectral Granger causality and transfer entropy, whose utility for BOLD time series has been questioned^60^. Finally, mutual information as an FC metric that also covers non-linear associations between BOLD time series has been suggested to be a promising alternative to correlational metrics^61^, but our results do not support its use to assess age-related neural and cognitive decline. This again corresponds to evidence in simulated data^60^ and is in line with theoretical considerations emphasizing that non-linear metrics of FC are not necessarily better than linear metrics, but rather address distinct aspects of the similarity between two signals^61^. In sum, our empirical examination supports the use of correlational and distance metrics in order to detect neural and cognitive decline related to age, while questioning the utility of partial correlation at least for summarizing FC scores across different regions.

### Variability across Datasets, Brain Regions, and Individuals

In addition to this general tendency towards the superiority of correlational and distance metrics, we obtained dataset-specific effects that have also been reported previously^18^. Specifically, when using dynamic time warping constrained by an Itakura parallelogram as FC metric, age-related neural decline was robustly identified in the MBB dataset, whereas no effect was observed in the HCP-Aging cohort. The utilized MRI scanners and the underlying scanning parameters differed noticeably between both projects, as the MBB protocol used a repetition time of 1400 ms with 657 timepoints, while the single HCP-Aging sequences had a repetition time of 800 ms with 478 timepoints. Given that especially the sequence length and repetition time have been shown to affect the sensitivity of FC metrics towards network alterations^60, 62^, it appears plausible that the FC metric of choice, to a certain degree, depends on the acquired EPI sequence. Hence, our findings indicate that methodological considerations regarding the scanning parameters need to be considered, but it remains to be determined which FC metric suits best under which scanning conditions in empirical data.

In addition to the impact of the scanning sequence, we also observed inter-individual variation between four patients with malignant tumors from the BTC dataset. Particularly, in the case of the first patient (Figure 5A), correlational and distance metrics were sensitive to white matter decline between regions close to the tumor. However, with regard to the other patients, this was not the case, as other FC metrics such as transfer entropy or kernel-based mutual information (2^nd^ patient, Figure 5B), phase coherence and dynamic time warping constrained by an Itakura parallelogram (3^rd^ patient, Figure 5C) or partial correlation and the frequency metrics (4^th^ patient, Figure 5D) mimicked the existing white matter decline in the ipsilateral compared to the contralateral hemisphere. While having been scanned under the same conditions, the four patients differ in terms of age, sex, tumor location, and tumor type (only 4^th^ patient). Consequently, these descriptive observations could suggest that the choice of the appropriate FC metric may depend on the individuaĺs characteristics or on the regions of interest.

### Embedding Current Findings into a Theoretical FC Framework

Our empirical findings can be embedded in the theoretical FC framework proposed by Reid et al.^25^. They categorize properties essential for mechanistic inferences from FC data into three types: theoretical, methodological, and confounding properties. Theoretical properties refer to the characteristics of the neural connections or pathways investigated (e.g., directionality between two neural assemblies). Methodological properties comprise all methodological approaches to assess these theoretical properties (e.g., type of FC metric). Confounding properties consist of any factors that may induce bias (e.g., motion artifacts). Reid et al.^25^ emphasize the need for empirical validations of methods used in FC research and suggest that future FC studies should comprehensively describe and discuss the three properties of the framework in the context of their particular research question to refine their conclusions. Our approach aimed for empirical validation of different FC metrics (methodological property) to explore neural deteriorations induced by age, malignant tumors, and schizophrenia (theoretical property). Importantly, our findings serve as an empirical endorsement of the FC framework proposed by Reid et al.^25^, as indicated by the following examples: First, using partial correlation as FC metric eliminates the confounding effect of unmeasured neural activity (confounding property), but proves less effective than other FC metrics that do not consider directness in detecting global neural decline (theoretical property). Second, the dataset-specific effects described above reveal that the selection of FC metrics (methodological property) to assess age-related neural decline (theoretical property) may depend on the parameters of the underlying scanning protocol (methodological property). Lastly, the findings in patients with malignant tumors show that the choice of FC metric (methodological property) may also depend on the type and location of the brain regions of the individual between which neural impairments are assessed (theoretical property). These examples demonstrate that no single FC assessment is superior; the optimal approach depends on the theoretical property of interest, other methodological properties than the choice of the FC metric itself, and relevant confounding properties.

### Brain Perfusion Measured by PCASL As a Neural Representation of Age-related Cognitive Decline

Lastly, beyond the major scope of this study, we observed that volumetric and perfusion-based measures in the hub regions gained from structural MRI and PCASL reflect neural and cognitive decline more accurately than structural connectivity and FC outcomes assessed by DWI and rs-fMRI, respectively. This is particularly interesting with respect to PCASL since this sequence is not yet commonly used in clinical neuroimaging research. According to our findings, cerebral blood flow in the hub regions assessed by PCASL decreases substantially with age and shows the strongest associations with age-related cognitive decline. Notably, cognitive impairments reflect a robust transdiagnostic phenomenon in various fields such as psychiatry^63^, but associations between cognitive functioning and the underlying neural processes are mostly small and often not reliably detected^26, 64^. The robust correlation between reductions in the cerebral blood flow of central brain hubs and age-related cognitive decline identified in the current work can serve as a promising basis for future clinical neuroimaging studies that aim to identify the neural underpinnings of cognitive deficits. Thus, our results may leverage the use of PCASL to investigate neural correlates of behavior in human neuroimaging.

### Study Limitations and Future Directions

Our study navigates through multiple FC metrices with room for enhancement, while maintaining a solid foundation for future exploration. First, our empirical validation of FC require knowledge about the ground truth of aberrant FC patterns in the brain^25^. We used a multimodal approach to demonstrate that the mean volume, perfusion, and structural connectivity in the hub regions decrease with age among different datasets, assuming that this decline should also manifest in lower FC between hubs (ground truth). Given the known link between deteriorations in brain volumes and structural connectivity with impairments in FC^14^, this assumption stands on biologically plausible ground. Nonetheless, the relationship between brain structure and function is not fully explicit due to complex multi-synaptic interactions ^65^. There may persist a small uncertainty if age-related decreases in FC between the hub regions truly reflect a stable and replicable ground truth. To refine this understanding, future examinations could combine empirical factors that induce strong neural impairments with model-based simulated data to bolster biologically plausible assumptions and to decrease the uncertainty about the ground truth.

Secondly, we conclude that selecting a proper FC metric depends on the acquired EPI sequence and the type and location of the connections of interest. While our data do not provide clear-cut guidelines for metric selection, future studies could evaluate how certain scanning parameters, such as the session length or repetition time, affect the utility of different FC metrics in capturing neural decline. This could involve leveraging respective biophysical models or simulating specific brain networks to test if distinct brain pathways require different FC metrics to evaluate the sensitivity of several FC metrics regarding known patterns of information flow within these networks.

Thirdly, our selection of 20 representative metrics from four domains, based on their previous application in fMRI research, opens the field for further inquiry time series^24^. Future investigations are encouraged to test the applicability of these yet-unutilized metrics.

Lastly, apart from the gradient analysis, we limited our analyses to the volume space to reduce the complexity of the current research approach, but future studies in this field may examine the use of different FC metrics in the vertex space.

## Conclusion

To conclude, we first provide empirical evidence that the utilized FC metric strongly affects the results in the context of typical fMRI research approaches. Secondly, our results demonstrate that correlational and distance metrics perform best in detecting neural decline, while questioning the utility of partial correlation when summarizing FC scores across regions are used. Thirdly, we demonstrate that the sensitivity of FC metrics towards neural decline is influenced by the parameters of the acquired EPI sequence, may vary between individuals that underwent the same scanning sequence, and depends on the regions of interest. Lastly, our results emphasize the promising role of the cerebral blood flow measured by PCASL as a neural representation of aging and cognitive impairment. These empirical findings strongly support the considerations by Reid et al.^25^, which illustrate the urgent need to define the respective theoretical, methodological, and confounding properties more carefully in future FC-based studies.

## Acknowledgements

Data collection and sharing for this project was provided by the Human Connectome Project (HCP; Principal Investigators: Bruce Rosen, M.D., Ph.D., Arthur W. Toga, Ph.D., Van J. Weeden, MD). HCP funding was provided by the National Institute of Dental and Craniofacial Research (NIDCR), the National Institute of Mental Health (NIMH), and the National Institute of Neurological Disorders and Stroke (NINDS). HCP data are disseminated by the Laboratory of Neuroimaging at the University of Southern California.

The data of the MBB project was obtained from the OpenNeuro database (DOI:10.18112/openneuro.ds000221.v1.0.0) with the accession number *ds000221*. The project was partially supported by the Volkswagen Foundation (AZ.: 89 440).

The data of the BTC project was obtained from the OpenNeuro database (DOI:10.18112/openneuro.ds001226.v4.0.0) with the accession number *ds001226*. This project was supported by the Special Research Funds (BOF) of the University of Ghent (01MR0210 and 01J10715), as well as Grant P7/11 from the Interuniversity Attraction Poles Program of the Belgian Federal Government.

The data of the CDP project was collected at the Department of Psychiatry and Psychotherapy of the LMU Hospital Munich and was supported by the BMBF with the EraNet project GDNF UpReg (01EW2206) to PF.

This work was endorsed by DZPG (German Center for Mental Health) to PF, FKZ: 01EE2303A, 01EE2303F.

## Authors’ contributions

LR and SW developed the initial research idea and performed all processing steps and analyses supervised by DK. LR and SW drafted the manuscript and designed the figures. LR, SW, and DK interpreted the results. DR created the formulas and descriptions of the different functional connectivity metrics. ZS proofed the statistical approaches and evaluated the description of the FC metrics. EW, FR, PF and AS provided data and assisted the planning of the project. DK and SS provided supervision and supported the development of further ideas for analyses. All authors discussed the results and revised the final manuscript.

## Availability of data, code, and material

Behavioral and imaging data of the HCP-Aging cohort are available through application at the Human Connectome Project, while data from the MBB and BTC cohort are freely available at OpenNeuro. Analysis commands, python scripts, and the singularity containers used will be published on a Github repository. Additional data can be made available upon request.

