## Supplemental Information for "How to measure functional connectivity using resting-state fMRI? A comprehensive empirical exploration of different connectivity metrics"

**Table S1**

Sample characteristics

| Data set | Group | N | Sex | Age (mean ± SD) |
|---|---|---|---|---|
| MBB | Healthy | 211 | 74 females<br>137 males | 144 young (20 – 35 years)*<br>67 old (55 – 80 years)* |
| CDP | Healthy | 149 | 76 females<br>72 males | 34.9 ± 12.5 |
| | Schizophrenia | 104 | 29 females<br>75 males | 38.5 ± 10.5 |
| HCP-Aging | Healthy | 719 | 403 females<br>316 males | 59.9 ± 15.8 |

Overview of the sample characteristics in all data sets. The sample sizes N refer to the maximum number of subjects that were considered in the statistical data analyses for the particular dataset. SD: standard deviation. *, ages were only available as categories in the MBB dataset.

**Table S2**. Scanning parameters.

| Dataset | Sequence | Resolution | TR | TE | FA | Slices | Duration | Acceleration factor | Gradient directions (b-value) |
|---|---|---|---|---|---|---|---|---|---|
| MBB | MP2-RAGE | 1.0 × 1.0 × 1.0 mm$^3$ | 5000 ms | 2.92 ms | 4/5° | 176 | 8min 22s | GRAPPA factor 3 | - |
| MBB | T2-weighted | 1.0 × 1.0 × 1.0 mm$^3$ | 3200 ms | 409 ms | var | 176 | 4 min 43s | GRAPPA factor 2 | - |
| MBB | EPI | 2.3 × 2.3 × 2.3 mm$^3$ | 1400 ms | 30 ms | 69° | 64 | 15min 30s | Multiband factor 4 | - |
| MBB | DWI | 1.7 × 1.7 × 1.7 mm$^3$ | 7000 ms | 80 ms | 90° | 88 | 9min 27s | Multiband factor 2 | 60 (1000 s/mm$^2$) |
| HCP-Aging | MP-RAGE | 0.8 × 0.8 × 0.8 mm$^3$ | 2500 ms | 2.2 ms | 8° | 208 | 8min 22s | GRAPPA factor 2 | - |
| HCP-Aging | T2 SPACE | 0.8 × 0.8 × 0.8 mm$^3$ | 3200 ms | 564 ms | 120 | 208 | 6min 35s | GRAPPA factor 2 | - |
| HCP-Aging | EPI | 2.0 × 2.0 × 2.0 mm$^3$ | 800 ms | 37 ms | 52° | 72 | 6min 40s | Multiband factor 8 | - |
| HCP-Aging | DWI | 1.5 × 1.5 × 1.5 mm$^3$ | 3230 ms | 89 ms | 78° | 92 | 5min 41s | Multiband factor 4 | 93 (3000 s/mm$^2$) |
| HCP-Aging | PCASL | 2.5×2.5×2.3 mm$^3$ | 3580 ms | 19 ms | 90° | 60 | 5min 29s | Factor 1 | - |
| CDP | MP-RAGE | 0.8 × 0.8 × 0.8 mm$^3$ | 2500 ms | 2.2 ms | 8° | 208 | 6min 54s | GRAPPA factor 2 | - |
| CDP | T2 SPACE | 0.8 × 0.8 × 0.8 mm$^3$ | 3200 ms | 564 ms | 120 | 208 | 6min 35s | GRAPPA factor 2 | - |
| CDP | EPI | 2.0 × 2.0 × 2.0 mm$^3$ | 800 ms | 37 ms | 52° | 72 | 6min 40s | Multiband factor 8 | - |
| BTC | MP-RAGE | 1.0 × 1.0 × 1.0 mm$^3$ | 1750 ms | 4.18 ms | 9° | 160 | 4min 05s | Factor 2 | - |
| BTC | EPI | 3.0 × 3.0 × 3.0 mm$^3$ | 2100 ms | 27 ms | 90° | 42 | 6min 24s | Factor 2 | - |
| BTC | DWI | 2.5×2.5×2.5 mm$^3$ | 8700 ms | 110 ms | 90° | 60 | 15min 14s | Factor 2 | 102 (2800 s/mm$^2$) |

Sequence, type of scanning sequence; resolution, voxel size; TR, Time of repetition; TE, echo time; TI, inversion time; FA, flip angle; slices, number of acquired slices; MP-RAGE, T1-weighted magnetization prepared rapid gradient echo; MP2-RAGE, T1-weighted magnetization prepared two rapid gradient echoes; EPI, echo planar imaging.

**Preprocessing of T1- and T2-weighted Images**

T1- and T2-weighted images of the MBB and HCP-Aging datasets were quality controlled with the automated quality control software MRIQC[1] and processed using the *recon-all* pipeline from FreeSurfer v7.2[2-4]. The pipeline included motion correction and averaging[5], removal of non-brain tissue[6], automated Talairach transformation, segmentation of the subcortical white matter and grey matter volumes[4, 7], intensity normalization[8], tessellation of the grey matter – white matter boundary, automated topology correction[9, 10] and surface deformation[2, 11, 12]. Further details on the FreeSurfer pipeline can be found under http://surfer.nmr.mgh.harvard.edu/. The segmentations were quality controlled based on visual inspection and erroneous cases were removed. Cortical and subcortical brain volumes were extracted from the FreeSurfer output using the *asegstats2table* function. Cortical regions were parcellated based on the Desikan-Killiany-Tourville (DKT)[13], while for the subcortical regions the default FreeSurfer parcellation was used. Volumes were corrected by the intracranial volume across all subjects that were included in the final analyses based on the proportions method[14].

**Preprocessing of structural DWI Images**

DWI images of the MBB, HCP-Aging, and BTC datasets were quality controlled before and after denoising and after preprocessing based on visual inspection and were further processed using functions from MRtrix3 v3.0.3 [15], FSL v6.0[16, 17], Freesurfer v7.2[2-4], AFNI v22.1.09[18, 19] and ANTS v2.3.5[20].

First, DWI images were denoised (MRtrix3: *dwidenoise*)[21], corrected for motion, eddy currents and susceptibility-induced distortions (MRtrix3: *dwifslpreproc*)[22, 23], and for bias field inhomogeneities (MRtrix3: *dwibiascorrect*, ANTS: *N4BiasFieldsCorrection*)[24]. Based on the *dhollander* algorithm, subject-specific response functions were estimated for each b-value and for three tissue types (MRtrix3: *dwi2response*). Fiber orientation densities were created based on multi-shell multi-tissue constrained spherical deconvolution (MRtrix3: *dwi2fod*)[25] and were intensity normalized (MRtrix3: *mtnormalise*). After increasing the intensity contrast (AFNI: *3dUnifize*), the T1w images were segmented into five tissue types (MRtrix3: *5ttgen*, FSL: *fast*)[26]. The averaged DWI images were co-registered to the gray matter segmentation (FSL: *flirt*)[27, 28], the resulting transformation matrix was used to linearly register the segmented T1w images to the diffusion space (MRtrix3: *mrtransform*) and tissue boundaries were created (MRtrix3: *5tt2gmwmi*). Anatomically constrained tractography was performed with ten million streamlines per subject (MRtrix3: *tckgen*)[29] and streamlines were refined using spherical deconvolution informed filtering of tractograms (MRtrix3: *tcksift2*)[30]. Structural connectivity between all regions from the subcortical parcellations from FreeSurfer and the cortical DKT atlas was estimated as the number of white matter tracts standardized by the size of the respective regions (MRtrix3: *tck2connectome*).

**Preprocessing of PCASL images**

PCASL images of the HCP-Aging dataset were processed using Oxford_ASL from FSL v6.0 as a command line interface of the Bayesian Inference for Arterial Spin Labeling MRI (BASIL) toolbox[31]. The pipeline includes a motion correction, a registration to the structural native space and to a standard template, a distortion correction, a calibration, and a partial volume correction[32]. Distortion correction was done based on respective fieldmaps gained from TOPUP[22], calibration was performed using a M0 calibration image, and partial volume correction was enabled for both grey and white matter based on segmentations from FAST[26]. Region analysis was performed based on regions from the subcortical FreeSurfer segmentations and the cortical DKT atlas.

**Preprocessing of functional EPI Images**

The resting-state functional EPI images of all datasets were quality controlled with MRIQC[1] and preprocessed using fMRIPrep v22.1.1[33] The following steps were executed: T1-weighted (T1w) volumes were corrected for INU (intensity non-uniformity) using *N4BiasFieldCorrection v2.1.0*[24] and skull-stripped using *antsBrainExtraction.sh v2.1.0* (OASIS template)[20]. Brain-extracted images were spatially normalized to the brain-extracted ICBM 152 Nonlinear Asymmetrical template version 2009c[34] using nonlinear registration within the *antsRegistration* tool of ANTs v2.1.0[35]. Brain tissue segmentation was performed using *fast* from FSL v5.0.9[16, 26]. Resting-state fMRI data was slice time corrected with *3dTshift* from AFNI v16.2.07[18] and motion corrected utilizing *mcflirt* from FSL v5.0.9[16, 28]. Distortion correction was performed by co-registering the fMRI image to the same-subject T1w image with intensity inversion[36, 37] constrained by an average fieldmap template[38] implemented in *antsRegistration*[35]. Thereafter, co-registration to the corresponding T1w image using boundary-based registration[39] with twelve degrees of freedom was executed with *flirt* from FSL v5.0.9[16, 27, 28]. Motion correcting transformations, field distortion correcting warp, BOLD-to-T1w transformation and T1w-to-template (MNI) warp were administered in one step using *antsApplyTransforms* from ANTs v2.1.0[20]. Frame-wise displacement (FD)[40] was computed for every functional run using the implementation in Nipype[41]. Automatic Removal Of Motion Artifacts based on independent component analysis (ICA-AROMA) was utilized to extract aggressive noise regressors[42]. For more details of the fMRIPrep pipeline see https://fmriprep.readthedocs.io/en/stable/workflows.html.

### Quality control procedures

Quality control of T1- and T2-weighted structural images and functional EPI images was done using the automated quality-control software MRIQC[1]. We computed different sequence-specific quality metrics, including the signal-to-noise ratio (SNR), contrast-to-noise ratio (CNR), coefficient of joint variation (CJV), entropy focused criterion (EFC), foreground-background energy ratio (FBER), median intensity non-uniformity (INU), Full Width at Half Maximum (FWHM), framewise displacement (FD), temporal SNR and the spatial root mean square after temporal differencing (DVARS). We documented subjects with low quality data in at least one of these image quality metrics. After structural image processing, we visually examined the resulting segmentations from FreeSurfer and excluded erroneous cases. After preprocessing and smoothing of functional images, we computed FD and temporal SNR and excluded images with $FD > 0.4$ or temporal $SNR < 80$. For both structural and functional images, we visually inspected the data distributions of volumes and FC, respectively, to identify outliers based on the 1.5 interquartile range cut-off. Outliers were not automatically excluded, but their image quality metrics were evaluated again. If an outlier showed clear deviations from the rest of sample in at least one of the abovementioned image quality metrics, it was excluded. The exclusion was performed prior to statistical data analysis.

For DWI and PCASL data, outliers in structural connectivity and brain perfusion, respectively, were identified based on the 1.5 interquartile range cut-off. Raw images and preprocessed images of these outliers were visually inspected and excluded in the case of relevant artifacts.

#### Hippocampal and medial frontal seeds

As part of the first research question, we examined the associations between cognitive functioning measured by the BACS in the CDP cohort and FC between hippocampal and medial frontal regions defined by the subcortical parcellations from FreeSurfer and the cortical DKT atlas. Table S3 summarizes all the regions whose connections we considered:

**Table S3**

Regions examined for links between hippocampal-frontal FC and cognition

| Abbreviation | Region |
|---|---|
| **Hippocampal seeds** | |
| L.PHIG | Left parahippocampal gyrus |
| R.PHIG | Right parahippocampal gyrus |
| L.EC | Left entorhinal cortex |
| R.EC | Right entorhinal cortex |
| L.HI | Left hippocampus |
| R.HI | Right hippocampus |
| **Middle frontal seeds** | |
| L.CACG | Left caudal anterior cingulate gyrus |
| R.CACG | Right caudal anterior cingulate gyrus |
| L.RACG | Left rostral anterior cingulate gyrus |
| R.RACG | Right rostral anterior cingulate gyrus |
| L.CMFG | Left caudal middle frontal gyrus |
| R.CMFG | Right caudal middle frontal gyrus |
| L.RMFG | Left rostral middle frontal gyrus |
| R.RMFG | Right rostral middle frontal gyrus |

List of regions from the subcortical FreeSurfer parcellations and the cortical DKT atlas used to examine the link between hippocampal-frontal FC and cognition across 20 FC metrics

**Metrics of functional connectivity**

20 FC metrics from four distinct categories (correlational, distance, frequency, and information-theoretic metrics) defined by Cliff et al.[43] were chosen. A metric was selected if its usage has already been demonstrated to be feasible in the context of fMRI research. Table S4 provides a categorical scheme that characterizes all metrics according to their mathematical properties. Respective formulas are described in the following paragraphs with a particular focus on BOLD timeseries extracted from rs-fMRI data:

<u>Pearson´s correlation</u>

The Pearson correlation coefficient measures linear relationship between two BOLD timeseries x and y and is defined as:

$$P_{X,Y} = \frac{Cov(x,y)}{\sigma_x \sigma_y} \tag{1}$$

with Cov(x,y) reflecting the covariance of both BOLD timeseries and $\sigma_x$ and $\sigma_y$ representing the standard deviations of x and y, respectively.

<u>Partial correlation</u>

Partial correlation assesses the linear relationship between two BOLD timeseries x and y, while considering the influence a third BOLD timeseries z. It is defined as:

$$\rho_{(x,y)/z} = \frac{P_{xy} - P_{xz} \cdot P_{yz}}{\sqrt{1 - P_{xz}^2}\ \sqrt{1 - P_{yz}^2}} \tag{2}$$

$P_{xy}$, $P_{xz}$, $P_{yz}$ are the Pearson´s correlation coefficients between BOLD timeseries x and y, x and z, and y and z, respectively.

<u>Spearman´s rho</u>

Spearman’s rank correlation coefficient assesses monotonic relationships between two BOLD timeseries x and y and is determined as the Pearson´s correlation coefficient between the rank variables R(x) and R(y). It is defined as:

$$r_s = P_{R(x),R(y)} = \frac{\mathrm{Cov}\big(R(x), R(y)\big)}{\sigma_{R(x)} \sigma_{R(y)}}. \tag{3}$$

Kendall´s tau

Kendall´s tau assesses monotonic relationships between two BOLD timeseries x and y and is defined as:

$$\tau_K = \frac{C - D}{C + D} \quad (4)$$

with C as the number of concordant pairs and D as the number of discordant pairs of timepoints. Any pair of observations ($x_i$, $y_i$) and ($x_j$, $y_j$), where i < j, is considered concordant if the sorting order of ($x_i$, $x_j$) and ($y_i$, $y_j$) is consistent. In other words: Both $x_i > x_j$ and $y_i > y_j$ or both $x_i < x_j$ and $y_i < y_j$ must hold for concordance.

Cross correlation

Cross correlation reflects the Pearson´s correlation between a defined number of time-shifted versions of the two BOLD timeseries x and y. It is defined as:

$$R_{x(t)y(t)}(\tau) = \frac{E\left[\left(x(t) - \mu_{x(t)}\right)\overline{\left(y(t+\tau) - \mu_{y(t)}\right)}\right]}{\sigma_{x(t)} \cdot \sigma_{y(t)}} \quad (5)$$

with the time difference denoted as τ (also known as displacement), the means $\mu_{x(t)}$ and $\mu_{y(t)}$ of the time-shifted BOLD timeseries and their standard deviations $\sigma_{x(t)}$ and $\sigma_{y(t)}$. $E\left[\left(x(t) - \mu_{x(t)}\right)\overline{\left(y(t+\tau) - \mu_{y(t)}\right)}\right]$ reflects the expected value and $\overline{\left(y(t+\tau) - \mu_{y(t)}\right)}$ is the respective complex conjugate.

Euclidean distance

The Euclidean distance between two BOLD timeseries x and y reflects the root of the sum of squared distances between corresponding timepoints of x and y in the Euclidean space. It is given by:

$$\mathrm{d}(x, y) = \sqrt{(x_1 - y_1)^2 + \cdots + (x_n - y_n)} \quad (6)$$

with $x_i$ and $y_i$ being the coordinates of the timepoints of both BOLD timeseries x and y located in the Euclidean space.

Cityblock distance

The Cityblock distance between two BOLD timeseries x and y is the sum of the absolute distances between corresponding timepoints $x_i$ and $y_i$ in the real coordinate space. It is defined by

$$d_C(x, y) = \sum_{i=1}^{n} |x_i - y_i| \quad (7)$$

Cosine distance

The Cosine Distance quantifies the dissimilarity between two BOLD timeseries x and y positioned to each other at an angle Θ. This distance measure is derived from the Euclidean dot product formula (represented by $*$) and is characterized by the equation:

$$d_{SC}(x, y) := cos(\Theta) = \frac{x * y}{||x|| \cdot ||y||} \tag{8}$$

Dynamic time warping

Dynamic Time Warping compares two BOLD timeseries x and y by finding the shortest distance between two random timepoints $x_i$ and $y_i$. Dynamic time warping is commonly employed with certain constraints for optimization purposes, such as the Itakura parallelogram and the Sakoe-Chiba band. The general formula is expressed as:

$$d_{DTW}(x, y) = \min_{\pi \in A(x,y)} \left\{ \sum_{i,j \in \pi} d(x_i, y_j) \right\} \tag{9}$$

where the alignment path π of length K is a sequence of K index pairs, A(x,y) reflects the set of all admissible paths and $d(x_i, y_i)$ represents the distance between two timepoints from the BOLD timeseries x and y.

In case of the Itakura parallelogram, the alignment path π is guided by a parallelogram, which limits the set of points from the sequences eligible for comparison. With regard to the Sakoe-Chiba band, the timepoint comparisons are constrained both horizontally and vertically within a specified corridor.

Coherence magnitude

Coherence magnitude captures the linear relationship of two BOLD timeseries x and y in the frequency domain and is given by:

$$C_{xy}(f) = \frac{|S_{xy}(f)|^2}{S_{xx}(f) S_{yy}(f)} \tag{10}$$

where $S_{xy}(f)$ is the cross-spectral density between x and y, while $S_{xx}(f)$ and $S_{yy}(f)$ denote the auto-spectral density of signals x and y, respectively.

Phase coherence

Coherence phase assesses temporal alignment or phase difference between two BOLD timeseries x and y. It is given by

$$\phi(f) = arg(S_{xy}(f)) \tag{11}$$

where arg(z) with a random complex number z indicates the angle formed between the positive real axis and the vector representing z in the complex plane. $S_{xy}(f)$ is the the cross-spectral density between the BOLD timeseries x and y.

Phase-locking value

The Phase Locking Value quantifies the phase consistency between two BOLD timeseries x and y and is defined by

$$\mathrm{PL}V_{xy} = \left|e^{i\phi_x(t)-\phi_y(t)}\right| \tag{12}$$

where $\phi_x(t) - \phi_y(t)$ is the phase difference between both timeseries and i the imaginary unit.

Phase-slope index

The Phase Slope Index quantifies the directional relationship between the phase of one BOLD timeseries x and the spatial gradient of phase of another brain region y. It is given by:

$$\psi_{xy} = \mathrm{Im}\left(\sum_{f\in F} \overline{C_{xy}(f)}\, C_{xy}(f+\delta f)\right) \tag{13}$$

where

$$C_{xy}(f) = \frac{S_{xy}(f)}{\sqrt{S_{xx}(f)S_{yy}(f)}} \tag{14}$$

is the complex coherency, Im denotes the imaginary part and F is the set of frequencies f over which the slope is summed.

Spectral Granger causality

Spectral Granger causality assesses the influence of one BOLD timeseries x on another BOLD timeseries y in the frequency domain. It is given by:

$$\mathrm{SG}C_{x\to y}(f) = \ln\left(\frac{S_{yy}(f)}{S_{yy}(f) - \left(\Sigma_{xx}\frac{\Sigma_{yx}^2}{\Sigma_{yy}}\right)\left|H_{yx}(f)\right|^2}\right) \tag{15}$$

where $S_{ij}$ with i and j being either x or y is the cross spectral densitiy matrix between a pair of timeseries x and y at a given frequency f. Σij is the covariance of the residuals of the autoregressive model and $H_{yx}(f)$ reflects the frequency-dependent spectral transfer matrix.

Mutual information

Mutual information measures the shared information between two BOLD timeseries x and y. It is a metric of the amount of information BOLD timeseries x contains about another BOLD timeseries y. It is defined as:

$$\mathrm{I}(x,y) = \int_x \int_y p(x',y') log\left(\frac{p(x',y')}{p(x')p(y')}\right) dxdy \tag{16}$$

with p(x',y') reflecting the joint probability mass function and p(x') and p(y') representing the marginal probability mass functions.

Mutual information with a Gaussian density estimation a Gaussian kernel centered at each data point is used to estimate the joint probability p(x',y'). Mutual information with a kernel-based density estimation a kernel function that applies weights to nearby data point is used to estimate the joint probability p(x',y').

Mutual information with a Kraskov-Stögbauer-Grassberger density estimation is based on the nearest neighbor distances. Particularly, the distances between x and y are considered by averaging the distance from each point to its k-nearest neighbor. It is defined as:

$$I_K(x,y) = \psi(k) + \psi(n) - \frac{\sum_{i=1}^{n}\left(\psi(k_i) - \psi(n-1)\right)}{n}. \tag{17}$$

with ψ() reflecting digamma functions, $k_i$ representing the amount of the neighbors and n the number of timepoints.

Transfer entropy

Transfer entropy quantifies the directional information flow between two BOLD timeseries x and y. In general, it is given by

$$\mathrm{T}(\mathrm{x} \rightarrow \mathrm{y}) = \mathrm{H}(y) - \mathrm{H}(y|x) \tag{18}$$

with H(y) and H(y|x) reflecting the entropies of respective timeseries. To compute the entropies an estimation of of the conditional distribution is required.

The transfer entropy with a Gaussian density estimation corresponds to the formula described in equation 19, whereby the entropies H(y) and H(y|x) are estimated using the conditional distribution p(x'|y'), which is determined by employing Gaussian Kernel density estimation. The transfer entropy with a Kraskov-Stögbauer-Grassberger density estimation also mirrors the equation outlined in 19. The entropies H(y) and H(y|x) are estimated via the conditional distribution p(x'|y'), utilizing the distances to the k-nearest neighbors in the joint space of x and y. Both versions can also be expressed as:

$$\mathrm{T}(\mathrm{x} \rightarrow \mathrm{y}) = \int p(y') \sum_{x} p(x'|y') log\left(\frac{p(x'|y')}{p(x')}\right) dy \tag{19}$$

**Table S4**

Categorization scheme of FC metrics

| Category | Metric | Type | Directionality | Directness | Domain | Linearity |
|---|---|---|---|---|---|---|
| Basic | Pearson´s correlation | similarity/dissimilarity | non-directional | non-direct | time | linearity assumption |
| | Partial correlation | similarity/dissimilarity | non-directional | direct | time | linearity assumption |
| | Spearman´s rho | similarity/dissimilarity | non-directional | direct | time | linearity assumption |
| | Kendall´s tau | similarity/dissimilarity | non-directional | direct | time | linearity assumption |
| | Cross correlation | similarity/dissimilarity | non-directional* | non-direct | time | linearity assumption |
| Distance | Euclidean distance | dissimilarity | non-directional | non-direct | time | no linearity assumption |
| | Cityblock distance | dissimilarity | non-directional | non-direct | time | no linearity assumption |
| | Cosine distance | dissimilarity | non-directional | non-direct | time | no linearity assumption |
| | Dynamic time warping (Itakura) | dissimilarity | non-directional | non-direct | time | no linearity assumption |
| | Dynamic time warping (Sakoe) | dissimilarity | non-directional | non-direct | time | no linearity assumption |
| Spectral | Coherence magnitude | similarity | non-directional | non-direct | frequency | linearity assumption |
| | Phase coherence | similarity | non-directional | non-direct | frequency | linearity assumption |
| | Phase-locking value | similarity | non-directional | non-direct | frequency | linearity assumption |
| | Phase-slope index | similarity | directional | non-direct | frequency | no linearity assumption |
| | Spectral Granger causality | similarity | directional | non-direct | frequency | linearity assumption |
| Information theoretic | Mutual information (Gaussian) | similarity | non-directional | non-direct | time | no linearity assumption |
| | Mutual information (kernel) | similarity | non-directional | non-direct | time | no linearity assumption |
| | Mutual information (Kraskov) | similarity | non-directional | non-direct | time | no linearity assumption |
| | Transfer entropy (Gaussian) | similarity | directional | non-direct | time | no linearity assumption |
| | Transfer entropy (Kraskov) | similarity | directional | non-direct | time | no linearity assumption |

Overview of the characteristics of the FC metrics examined in this study. The type indicates if the resulting value of metric provides information on similarity, dissimilarity, or both. Directionality indicates if a directional relationship between two timeseries is investigated. Directness describes if the metric adjusts for the influence of other regional timeseries than the two of interest. Domain specifies if the metric quantifies similarity in the time or frequency domain. Linearity indicates if the metric measures a linear relationship between two timeseries or not. * Cross correlation may cover directionality indirectly due to the time-shifting.

**Table S5**

Cognitive and Neuroimaging Outcomes per research aim and dataset

| Research Aim | Neuroimaging Outcome | Modality | Cognitive Outcome | Dataset | Reason for Particular Outcome |
|---|---|---|---|---|---|
| Investigate impact of the selected FC metric on several common FC-based approaches in rs-fMRI research (1st aim) | FC within default-mode network for each FC metric | Resting-state fMRI | - | MBB | Default-mode network connectivity as one common neural target in FC research |
| | Composition of gradients for each FC metric | Resting-state fMRI | - | MBB | Macroscale gradients as a common approach to describe the organization of the brain in FC research |
| | Hippocampal-frontal FC for each FC metric | Resting-state fMRI | BACS | CDP | Brain-behavior associations as a common approach in FC research. Hippocampal-frontal FC and cognition serves as a prominent example. |
| | FC for each FC metric between seven Yeo brain networks | Resting-state fMRI | - | CDP | FC-based case-control comparisons to identify disorder-related biomarkers as a common approach in FC research |
| Investigate which FC metric most accurately captures biologically plausible disruptions of neural connections (2nd aim) | Mean volume in hub regions | T1- and T2-weighted MRI | NIH Toolbox<br><br>- | HCP-Ageing<br><br>MBB | Age-related decline in regional volumes and associated cognitive impairments as a biologically plausible process that also impairs FC between respective regions |
| | Mean structural connectivity between hub regions | DWI | NIH Toolbox<br><br>- | HCP-Ageing<br><br>MBB | Age-related decline in white matter connectivity and associated cognitive impairments as a biologically plausible process that also impairs FC between respective regions |
| | Mean cerebral blood flow in hub regions | PCASL | NIH Toolbox | HCP-Ageing | Age-related decline in cerebral blood flow and associated cognitive impairments as a biologically plausible process that also impairs FC between respective regions |

| | | | | | |
|---|---|---|---|---|---|
| | Mean functional connectivity between hub regions | Resting-state fMRI | NIH Toolbox<br><br>- | HCP-Ageing<br><br>MBB | To examine how well different FC metrics reveal age-related neural deteriorations in hub regions and associated cognitive impairments |
| | Mean structural connectivity between regions close to malignant tumor | DWI | - | BTC | Disruptions of white matter tracts between regions close to the tumor location as a biologically plausible process that also impairs FC between respective regions |
| | Mean functional connectivity between regions close to malignant tumor | Resting-state fMRI | - | BTC | To examine how well different FC metrics reveal neural deteriorations in regions close to the malignant tumor |

Overview of all neural and cognitive outcomes examined for each research and dataset.

**Figure S1**

Absolute connectivity strength within the DMN of the different FC metrics

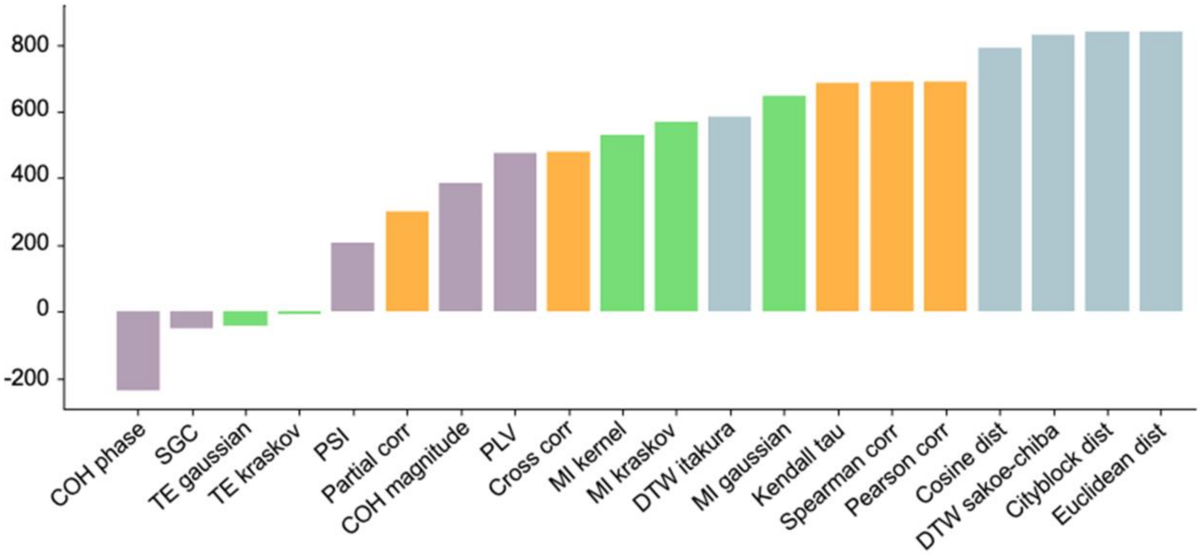

This figure displays the absolute connectivity strength within the DMN across the different FC metrics. The connectivity values within DMN regions are aggregated. The metrics are sorted based on the connectivity strength among the brain regions constituting the DMN. The y-axis represents the absolute values of the DMN for both hemispheres. The x-axis indicates the different metrics. Pearson corr, Pearson´s correlation; Partial corr, partial correlation; Spearman corr, Spearman´s correlation; Kendall tau, Kendall´s tau; Cross corr, cross correlation; Euclidean dist, Euclidean distance; Cityblock dist, Cityblock distance; Cosine dist, Cosine distance; DTW itakura, dynamic time warping constrained with Itakura parallelogram; DTW sakoe-chiba; dynamic time warping constrained with Sakoe-Chiba band; COH magnitude, coherence magnitude; COH phase, phase coherence; PLV, phase-locking value; PSI, phase slope index; SGC, spectral Granger causality; MI gaussian, mutual information with gaussian density estimation; MI kernel, mutual information with kernel-based density estimation; MI kraskov, mutual information with Kraskov-Stögbauer-Grassberger density estimation; TE gaussian, transfer entropy with gaussian density estimation; TE kraskov transfer entropy with Kraskov-Stögbauer-Grassberger density estimation.

**Gradient analysis**

As indicated in the results section of the main manuscript and in Figure S2, the direction and composition of the first and second gradient vary depending on the chosen FC metric. The original study by Margulies and colleagues[44] suggests that the first gradient originates from primary and unimodal visual, somatosensory/motor, and auditory regions, extending to regions such as the angular gyrus, rostral anterior cingulate, posteromedial cortex, middle temporal gyrus, and middle and superior frontal gyri, collectively forming the DMN. On the opposite end of the spectrum, the second encompasses occipital cortex regions involved in visual processing, with the other end comprising somatosensory and motor regions around the central sulcus, along with auditory regions of the temporal perisylvian region. For most FC metrics, we observed consistent patterns in the first and second gradients. The reversal of directions in these gradients suggests that a greater variance in BOLD timeseries is accounted for by either the DMN or its constituent regions. Despite utilizing resting-state data, the variance in BOLD timeseries within or associated with the DMN appears to be interchangeable.

**Figure S2**

Example illustration of the first and second gradient based on different FC metrics

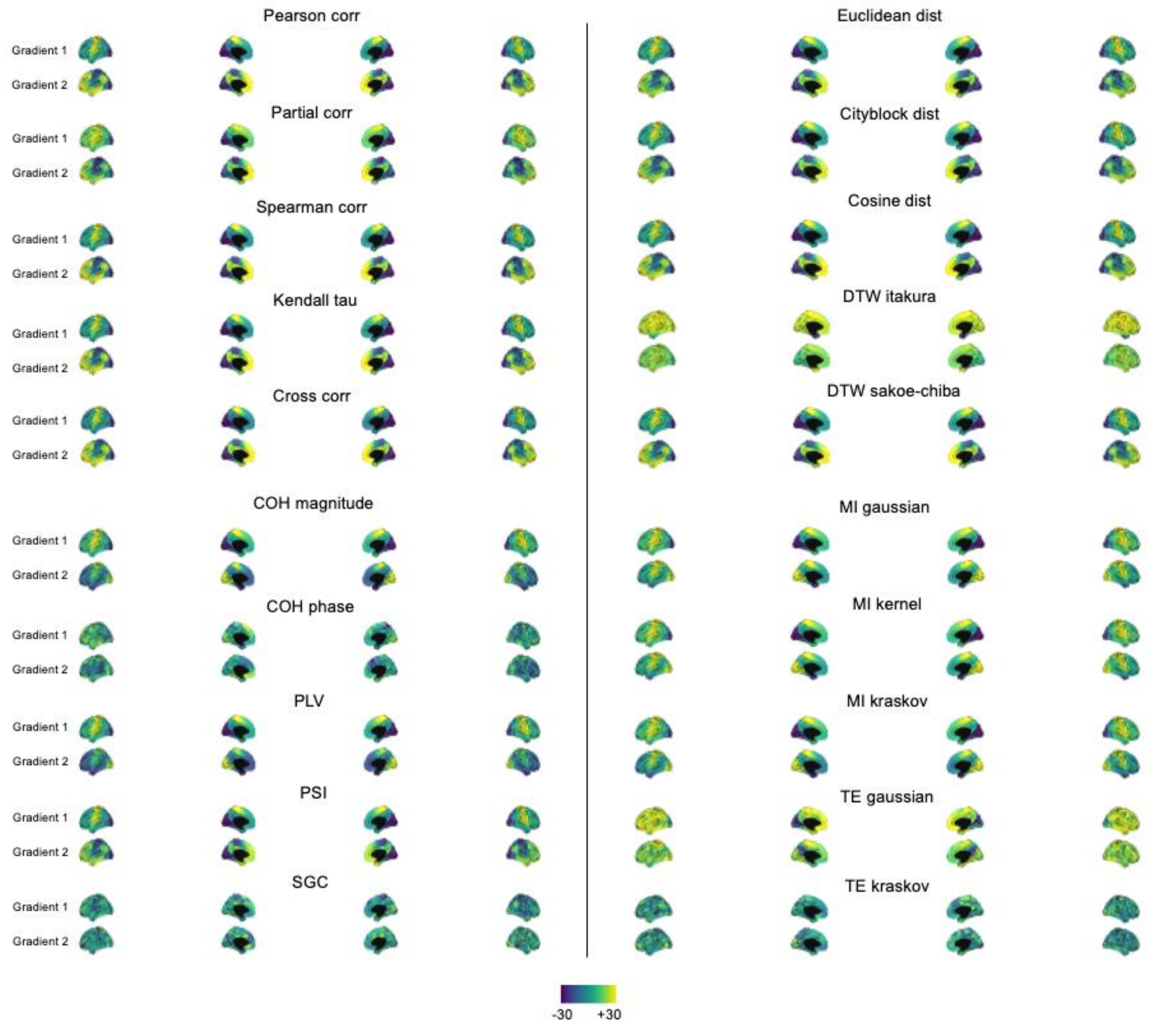

This figure depicts the primary and secondary gradients of FC metrics, which reflect the variance in the BOLD signal across different brain regions, indicating the levels of activity and strength of functional connectivity among them. Each gradient exhibits values ranging from negative to positive throughout the brain. The primary gradient captures spatial variations in brain activity, mapping regions involved in visual processing (depicted in blue) to the somatosensory network (depicted in yellow). The secondary gradient extends from primary and unimodal visual and somatosensory/motor regions (depicted in blue) to the DMN (depicted in yellow). The inverse of the second gradient of COH magnitude, PLV, MI Gaussian, MI kernel, and MI Kraskov manifests a directional trend from the DMN towards the visual and somatosensory/motor regions. However, COH phase, SGC, TE kraskov, and DTW itakura do not exhibit significant patterns related to visual, somatosensory network and DMN. Pearson corr, Pearson´s correlation; Partial corr, partial correlation; Spearman corr, Spearman´s correlation; Kendall tau, Kendall´s tau; Cross corr, cross correlation; Euclidean dist, Euclidean distance; Cityblock dist, Cityblock distance; Cosine dist, Cosine distance; DTW itakura, dynamic time warping constrained with Itakura parallelogram; DTW sakoe-chiba; dynamic time warping constrained with Sakoe-Chiba band; COH magnitude, coherence magnitude; COH phase, phase coherence; PLV, phase-locking value; PSI, phase slope index; SGC, spectral Granger causality; MI gaussian, mutual information with gaussian density estimation; MI kernel, mutual information with kernel-based density estimation; MI kraskov, mutual information with Kraskov-Stögbauer-Grassberger density estimation; TE gaussian, transfer entropy with gaussian density estimation; TE kraskov transfer entropy with Kraskov-Stögbauer-Grassberger density estimation.
